# Murine breast cancers disorganize the liver transcriptome in zonated manners

**DOI:** 10.1101/2022.09.27.509354

**Authors:** Alexis Vandenbon, Rin Mizuno, Riyo Konishi, Masaya Onishi, Kyoko Masuda, Yuka Kobayashi, Hiroshi Kawamoto, Ayako Suzuki, Masahito Shimizu, Yasuhito Tanaka, Yutaka Suzuki, Shinpei Kawaoka

## Abstract

The spatially organized gene expression program within the liver specifies hepatocyte functions according to their relative distances to the bloodstream (i.e., zonation), contributing to liver homeostasis. Despite the knowledge that solid cancers remotely disrupt liver homeostasis, it remains unexplored whether solid cancers affect liver zonation. Here, using spatial transcriptomics, we thoroughly investigate the abundance and zonation of hepatic genes in cancer-bearing mice. We find that breast cancers affect liver zonation in various distinct manners depending on biological pathways. Aspartate metabolism and triglyceride catabolic processes retain relatively intact zonation patterns, but the zonation of xenobiotic catabolic process genes exhibits a strong disruption. The acute phase response is induced in zonated manners. Furthermore, we demonstrate that breast cancers activate innate immune cells in particular neutrophils in distinct zonated manners, rather than in a uniform fashion within the liver. Collectively, breast cancers disorganize hepatic transcriptomes in zonated manners, thereby disrupting zonated functions of the liver.

## Introduction

Remote cancers affect the liver in various manners thereby disrupting host homeostasis^1–11^. For example, circadian gene expression rhythm is disrupted in a breast cancer model and a genetically induced lung cancer model^3,4^. This lung cancer model also disrupts hepatic glucose metabolism^9^. Fearon and colleagues showed that interleukin-6 derived from pancreatic ductal adenocarcinoma suppresses hepatic ketogenesis^2^. A zebrafish intestinal tumor reduces bile production in the liver^5^. We recently found that remote solid cancers dampen hepatic nitrogen metabolism^8^. Some of these abnormalities are detectable in human cancer patients^12,13^. These metabolic abnormalities in the liver are often accompanied by systemic phenotypes such as reduced behavioral activity, suggesting the significance of liver alteration in systemic phenotypes caused by remote cancers^1,8^. Of note, the above-described discoveries were made through analyses against whole liver tissues.

It has been unknown whether remote cancers disrupt spatially organized gene expression (i.e., zonated gene expression) in the liver. The liver has a spatially organized tissue structure known as zonation, formed by repetitive hexagonal units called liver lobules^14,15^. The lobules are associated with portal veins and central veins. Portal veins are at the junction of neighboring lobules, supplying nutrient- and oxygen-rich blood to nearby hepatocytes. Those hepatocytes eventually consume nutrients and oxygen. Consistent with this, hepatocytes nearby portal veins express genes critical for active energy metabolism. The consequently exhausted blood is then drained by central veins. In contrast to portal vein-associated hepatocytes, hepatocytes nearby central veins are enriched for xenobiotic metabolism genes. Thus, hepatocytes nearby portal veins and central veins are transcriptionally distinct^14^. Recent advances in spatial transcriptomics enable us to obtain zonated gene expression profiles from the liver both in physiological and disease conditions^16–25^. However, it remains unclear whether cancer-induced liver abnormalities are associated with the disruption in liver zonation.

Here, we use spatial transcriptomics to understand how remote cancers affect liver zonation. Combining spatial transcriptomics and single-cell RNA sequencing (scRNA-seq), we characterize the livers of breast cancer-bearing animals at the spatial and single-cell resolution, finding that murine breast cancers disrupt liver zonation. This study will be an important basis to understand how breast cancers affect spatially organized gene expression in the liver to disturb the whole liver homeostasis.

## Results

### Visualizing liver zonation using spatial transcriptome

To understand how cancers affect liver zonation, we performed spatial transcriptomic analyses against the livers of sham-operated and 4T1 cancer-bearing animals. We exploited this 4T1 model because we previously showed that 4T1 breast cancers remotely affect gene expression and metabolism in the liver^4,8^.

We first confirmed the previously reported zonated expression patterns of *Albumin (Alb)* and *Cyp2e1* in the livers of sham-operated mice (Fig. 1a and Fig. S1a-b). It has been known that hepatocytes nearby portal veins are supplied with nutrient- and oxygen-rich blood, and abundantly express *Alb*. On the other hand, hepatocytes nearby central veins express detoxifying genes including *Cyp2e1*. Exploratory analysis of spatially variable genes (SVGs) using singleCellHaystack^26^ confirmed two subsets of genes with the known expression patterns of *Alb* and *Cyp2e1*, demonstrating that these two genes are expressed in different zones in the liver (Fig. 1a).

**Figure 1:**
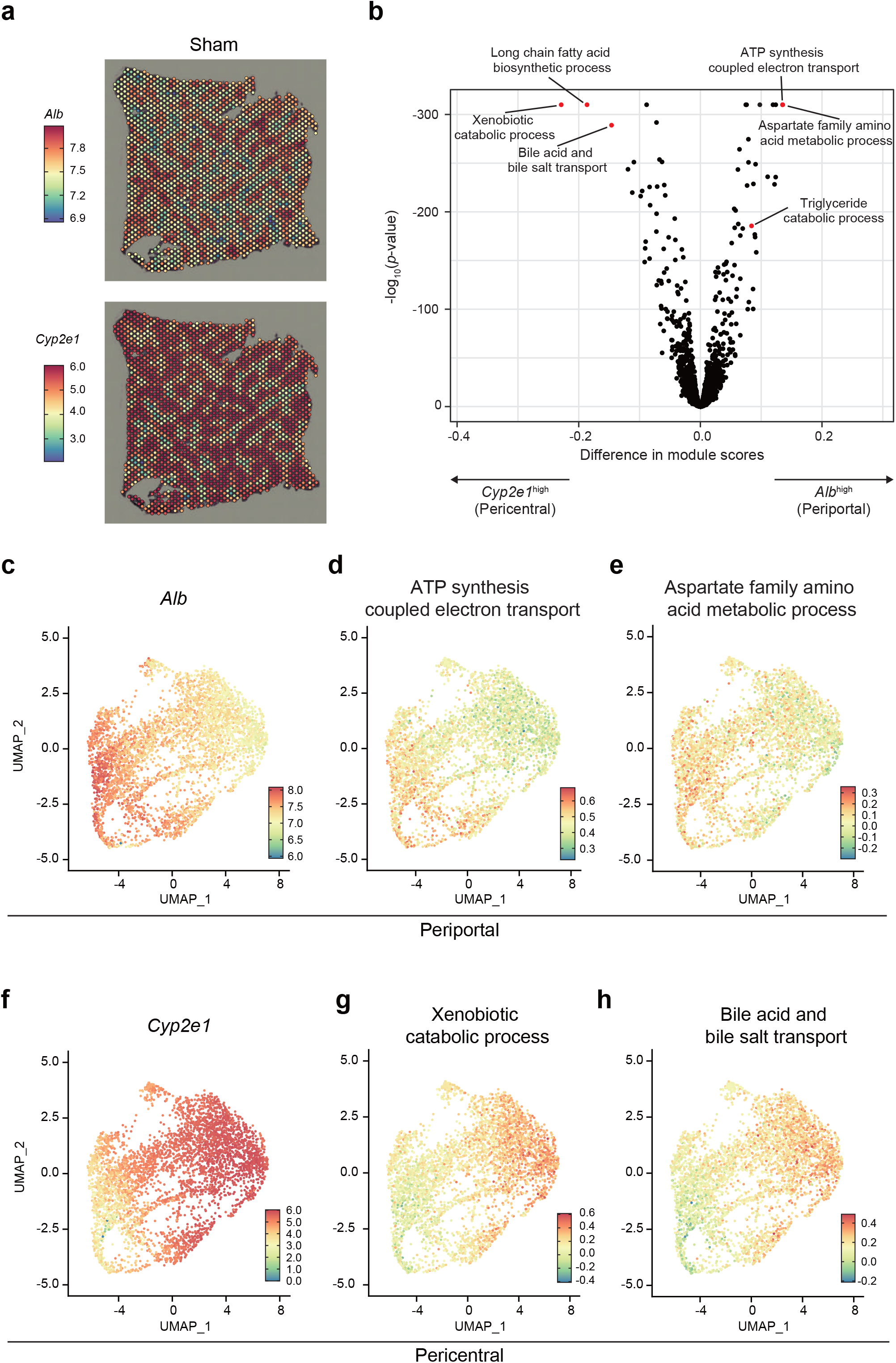
Visualization of liver zonation using spatial transcriptomics. **(a)** *Alb* and *Cyp2e1* expression in one of the sham samples. A second sample is shown in Fig. S1. **(b)** Volcano plot of the module scores of 2,898 GO terms. The x-axis shows the difference in module scores between *Alb*^high^ and *Cyp2e1*^high^ zones. The y-axis shows the p-values (-log1o) of a Wilcoxon Rank Sum test. Fig. S2 shows the same plot with more GO terms indicated. **(c-h)** UMAP plots of the Visium spots of the two sham samples. **(c)** UMAP_1 is anti-correlated with *Alb* expression. Spots on the left side represent the periportal zone with high *Alb* expression. Genes involved in. the ATP synthesis coupled electron transport **(d)** and the aspartate family amino acid metabolism **(e)** have high activity in the periportal zone. **(f)** UMAP_1 is correlated with *Cyp2e1* expression. Spots on the right side represent the pericentral zone with high *Cyp2e1* expression. Genes involved in the xenobiotic catabolic process **(g)** and the bile acid and bile salt transport **(h)** have high activity in the pericentral zone.

To further characterize these zonation patterns, we divided the Visium spots of each image into 3 sets of equal size: those with the highest *Alb* expression (*Alb*^high^), those with the highest *Cyp2e1* expression (*Cyp2e1*^high^), and the remaining one-third (with intermediate *Alb* and *Cyp2e1* expression). We then calculated “module scores” (average expression levels) using Seurat for sets of genes associated with 2,898 Gene Ontology (GO) terms, and compared the *Alb*^high^ and the *Cyp2e1*^high^ zones of the two sham samples (Fig. 1b and Fig. S2; Table S1). As shown in Fig. 1b, the *Alb*^high^ zone had higher expression of genes involved in “ATP synthesis coupled electron transport” and “aspartate family amino acid metabolic process.” This observation was further confirmed by the uniform manifold approximation and projection (UMAP) plots where the zonated expression of *Alb* was highlighted (Fig. 1c-e). In contrast, the *Cyp2e1*^high^ zone had higher expression of genes related to the “xenobiotic catabolic process” and “bile acid and bile salt transport” (Fig. 1b and Fig. 1f-h). Thus, our data confirm that different liver zones conduct distinct metabolic pathways.

### Breast cancers alter spatial transcriptomics in the liver

We next compared the spatial transcriptome of the livers of sham-operated and cancer-bearing mice in various manners. We found that both sham and cancer-bearing samples retained the zonated expression patterns of *Alb* and *Cyp2e1* (Fig. 2a and Fig. S1). This data suggested that distal cancers did not completely abolish zonated gene expression in the liver. However, when we analyzed the datasets via dimensionality reduction using principal component analysis (PCA) and UMAP, sham and cancer-bearing spots appeared nearly completely separated, even after batch effect correction (Fig. 2b, UMAP_2 axis, and Fig. S3). This lack of overlap between sham and cancer-bearing samples indicated strong changes in the cell composition and/or gene expression at each spot. In addition, this plot also exhibited a clear separation of *Alb^high^* versus *Cyp2e1^high^* spots along the UMAP_1 axis, further confirming the zonated gene expression in the liver in our datasets (Fig. 2b).

**Figure 2:**
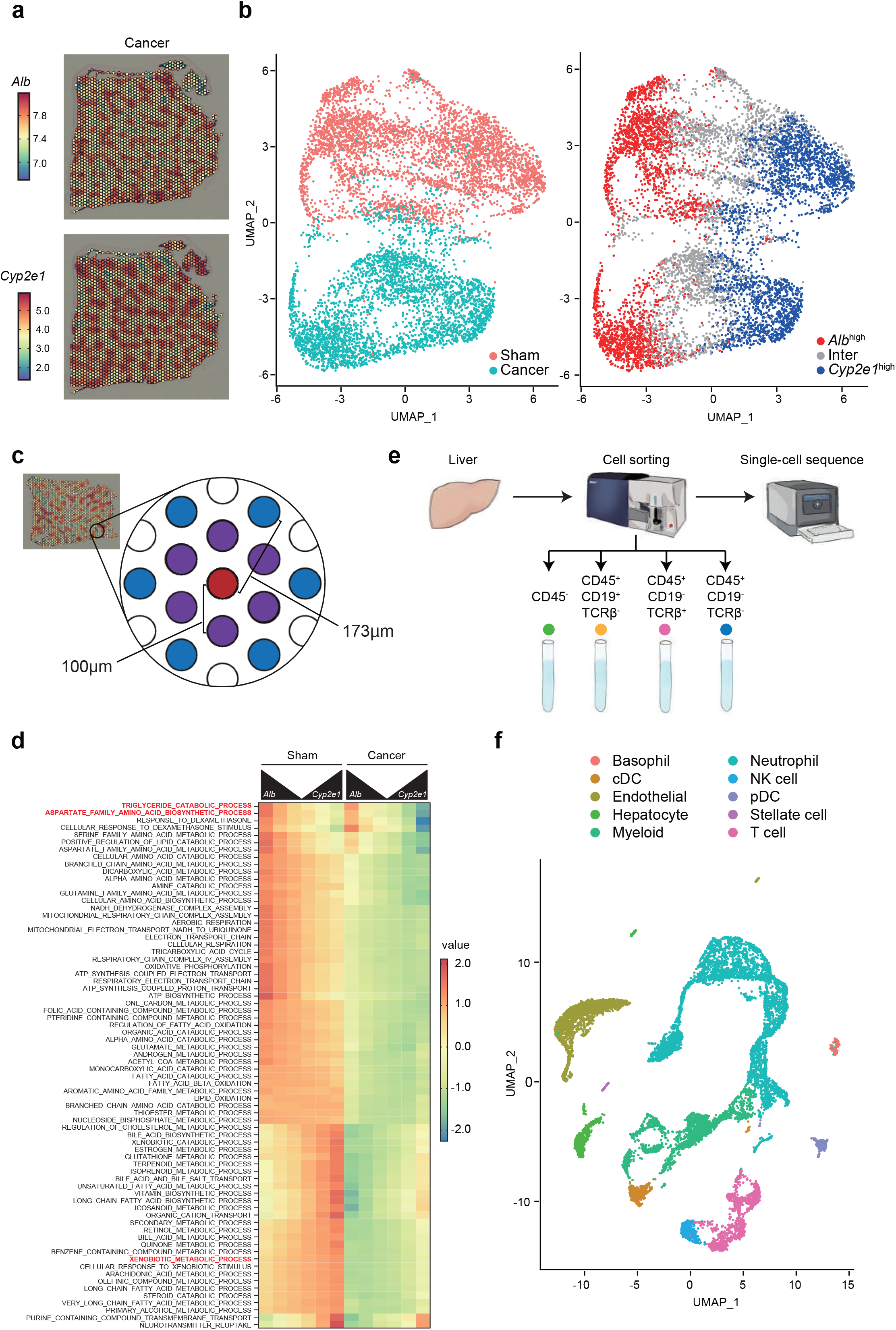
Spatial and single-cell characterization of the livers of cancer-bearing mice. **(a)** *Alb* and *Cyp2e1* expression in one of the 4T1 cancer samples. A second sample is shown in Fig. S1. **(b)** UMAP plot of the Visium spots of the two sham and two cancer-bearing samples. Left: spots are colored by condition, revealing a clear distinction between sham and cancer-bearing samples. Right: colors reflect the classification of spots according to *Alb* and *Cyp2e1* expression. **(c)** Illustration of the definition of closest (purple) and secondary (blue) neighboring spots of a spot of interest (red), and the approximate distances separating them. **(d)** Heatmap showing the normalized module scores of selected GO terms in function of their distance to *Alb*^high^ and *Cyp2e1*^high^ spots in sham (left) and cancer-bearing (right) samples. **(e)** The scheme for single-cell transcriptome. We sorted CD45^-^ cells, CD45^+^CD19^+^TCRβ^-^ cells, CD45^+^CD19^-^TCRβ^+^ cells, and CD45^+^CD19^-^TCRβ^-^ cells, mixing them at the same proportion among the samples. The obtained mixture of cells is subjected to single-cell transcriptomics. **(f)** UMAP plot of the scRNA-seq dataset, indicating cell type annotations. Fig. S7 shows the same data separated by condition.

To comprehensively investigate whether cancer transplantation affects liver zonation, we visualized the module scores of genes involved in distinct biological pathways according to the zonated expression of *Alb* and *Cyp2e1* (Fig. 2c, d and Table S2). In the analysis, we defined *Alb*^high^ spots to be the top 10% spots with the highest *Alb* expression in each image (the red spot in Fig. 2c). We then calculated mean module scores in these *Alb*^high^ spots and their closest (the 6 spots at a distance of 100 μm center-to-center; the purple spots in Fig. 2c) and secondary (the 6 spots at a distance of about 173 μm center-to-center; the blue spots in Fig. 2c) neighboring spots. We did the same thing for the top 10% spots with the highest *Cyp2e1* expression (*Cyp2e1*^high^ spots). This method validated the zonated gene expression of various metabolic pathways, for example, the “xenobiotic catabolic process” and the “aspartate family amino acid metabolic process” in the liver (Fig. 2d).

We added one more important layer to our analyses. As described earlier, the liver consists of multiple cell types. Given the size of the Visium spots (diameter of 55 μm), it is reasonable to expect that RNA molecules captured at each spot originated from >10 different cells. To address this issue, we performed single-cell transcriptomics analyses (Fig. 2e). We dissociated the whole liver tissues with collagenase and collected CD45^+^CD19^+^TCRβ^-^ B cells, CD45^+^CD19^-^TCRβ^+^ T cells, CD45^+^CD19^-^TCRβ^-^ immune cells that are neither B cells nor T cells, and other cell types in the liver (CD45^-^) including hepatocytes using flow cytometry. The sorted cells were mixed at the same proportion among different biological replicates and subjected to single-cell transcriptomic analysis. We determined the transcriptome of 11,085 cells from 4 biological replicates (*n*=2 for both the sham-treated group and 4T1 breast cancer-bearing group, respectively). After processing the data, we found 23 clusters of cells (Fig. S4a). Using known marker genes, we identified hepatocytes, hepatic stellate cells, endothelial cells, conventional dendritic cells, plasmacytoid dendritic cells, T cells, B cells, macrophages (including Kupffer cells), neutrophils, basophils, natural killer cells, and other monocyte-derived cells (Fig. 2f and Fig. S4b; Table S3). These scRNA-seq datasets allowed us to link the differentially expressed biological pathways with their underlying cell types, helping interpretation of our spatial transcriptomics datasets.

Using these datasets, we examined which cell types accounted for the changes in spatial transcriptomics. Such comprehensive analyses resulted in the following findings: (i) breast cancers rewire zonated gene expression patterns in hepatocytes (Fig. 3–4) and (ii) breast cancers activate innate immune cells particularly neutrophils in distinct zonated manners (Fig. 5). These findings are discussed in the following two sections.

**Figure 3:**
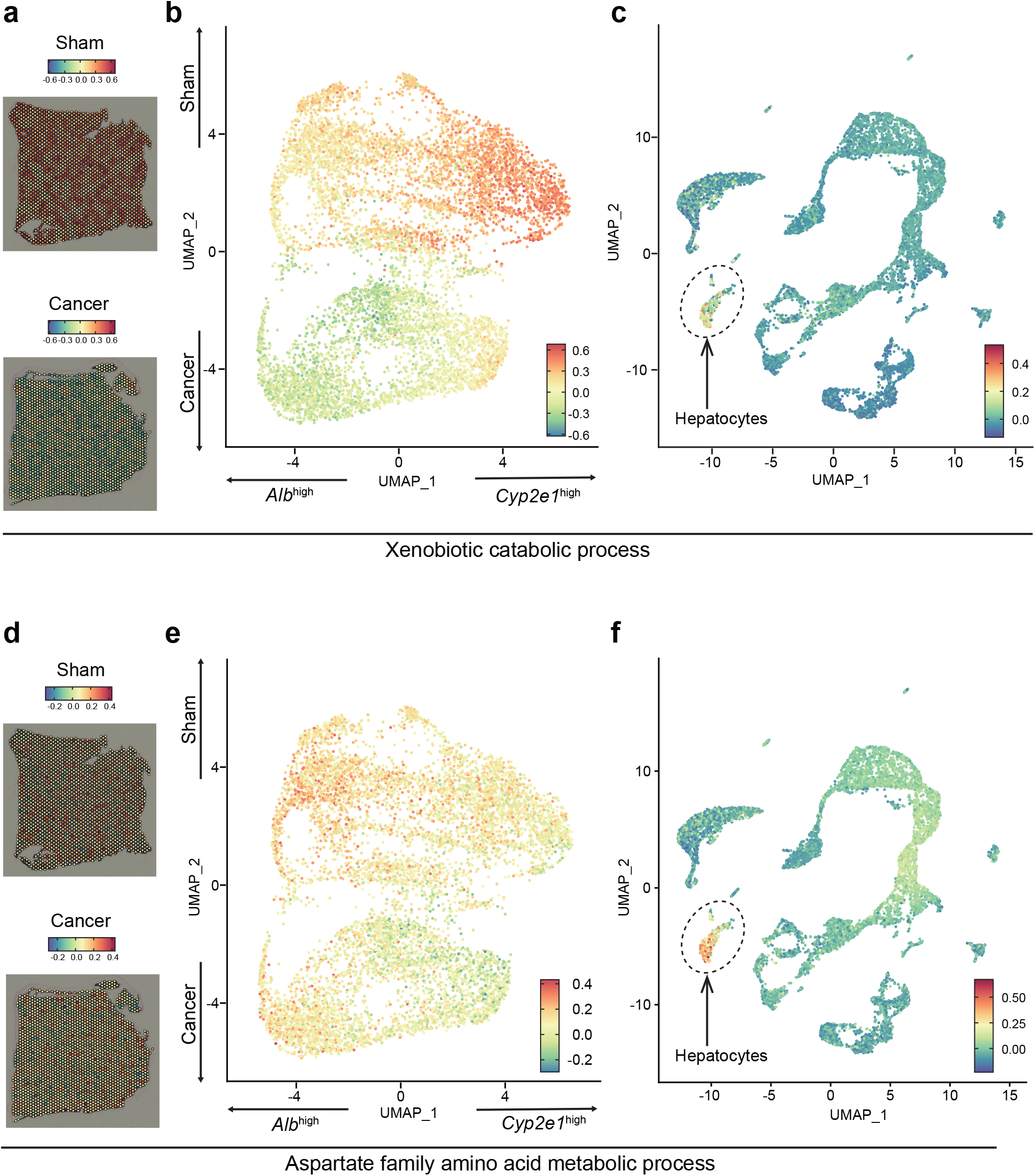
4T1 breast cancers disrupt liver zonation in various manners. **(a-c)** For the xenobiotic catabolic process. **(a)** Module scores in one of the sham and cancer Visium samples. **(b)** The same module scores in a UMAP representation of the Visium data. **(c)** Module scores in the scRNA-seq data, showing high scores predominantly in the hepatocyte cluster. **(d-f)** Same as **(a-c)** for genes associated with the aspartate family amino acid metabolic pathway.

**Figure 4:**
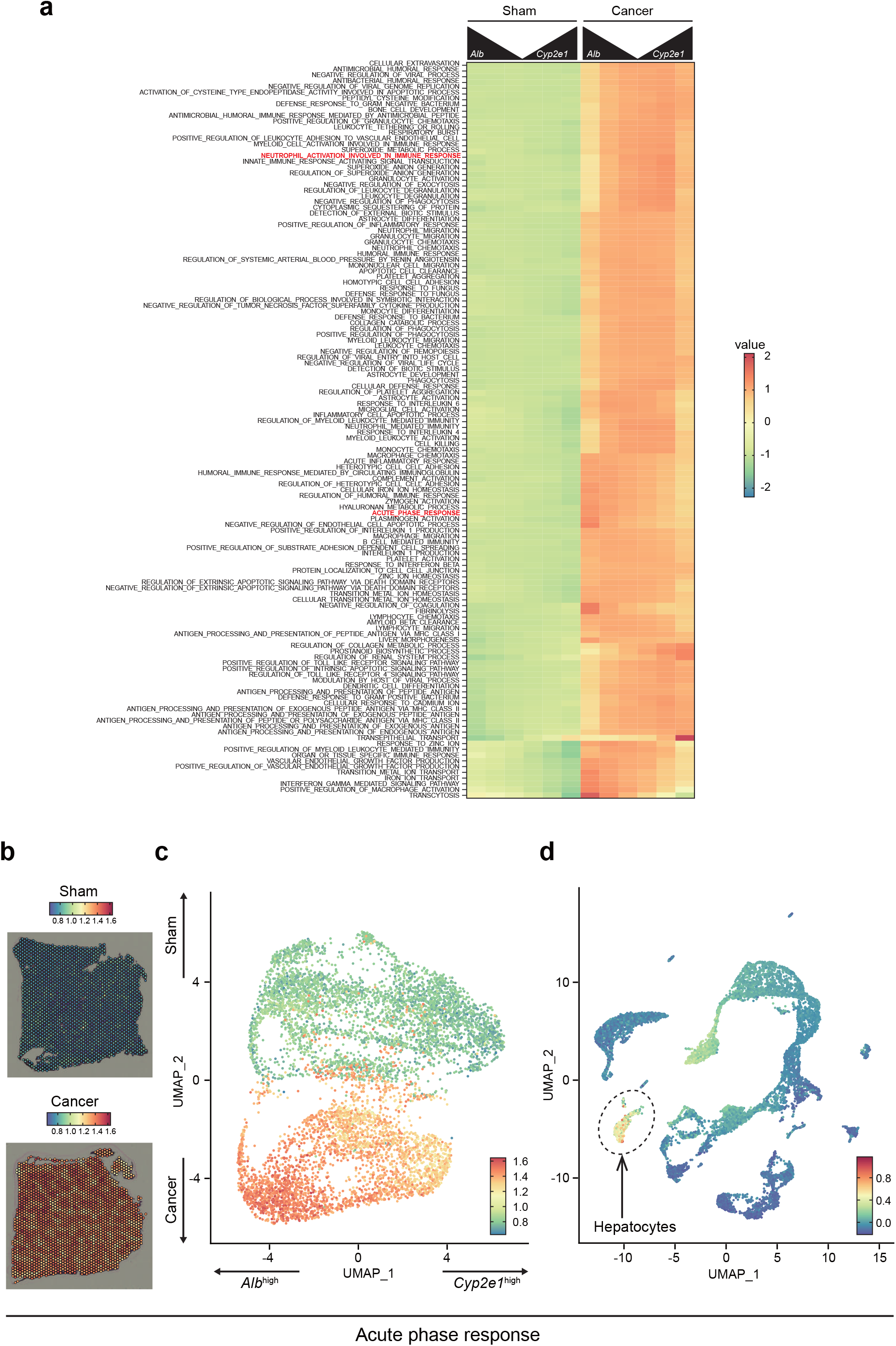
4T1 breast cancers induce acute phase response in a zonated manner. **(a)** Heatmap showing the normalized module scores of selected GO terms in function of their distance to *Alb*^high^ and *Cyp2e1*^high^ spots in sham (left) and cancer-bearing (right) samples. **(b-c)** Genes involved in the acute phase response are induced in the liver of cancer-bearing mice. **(b)** Module scores in one of the sham and cancer Visium samples. **(c)** The same module scores in a UMAP representation of the Visium data. **(d)** Module scores in the scRNA-seq data, showing high scores predominantly in the hepatocyte cluster.

**Figure 5:**
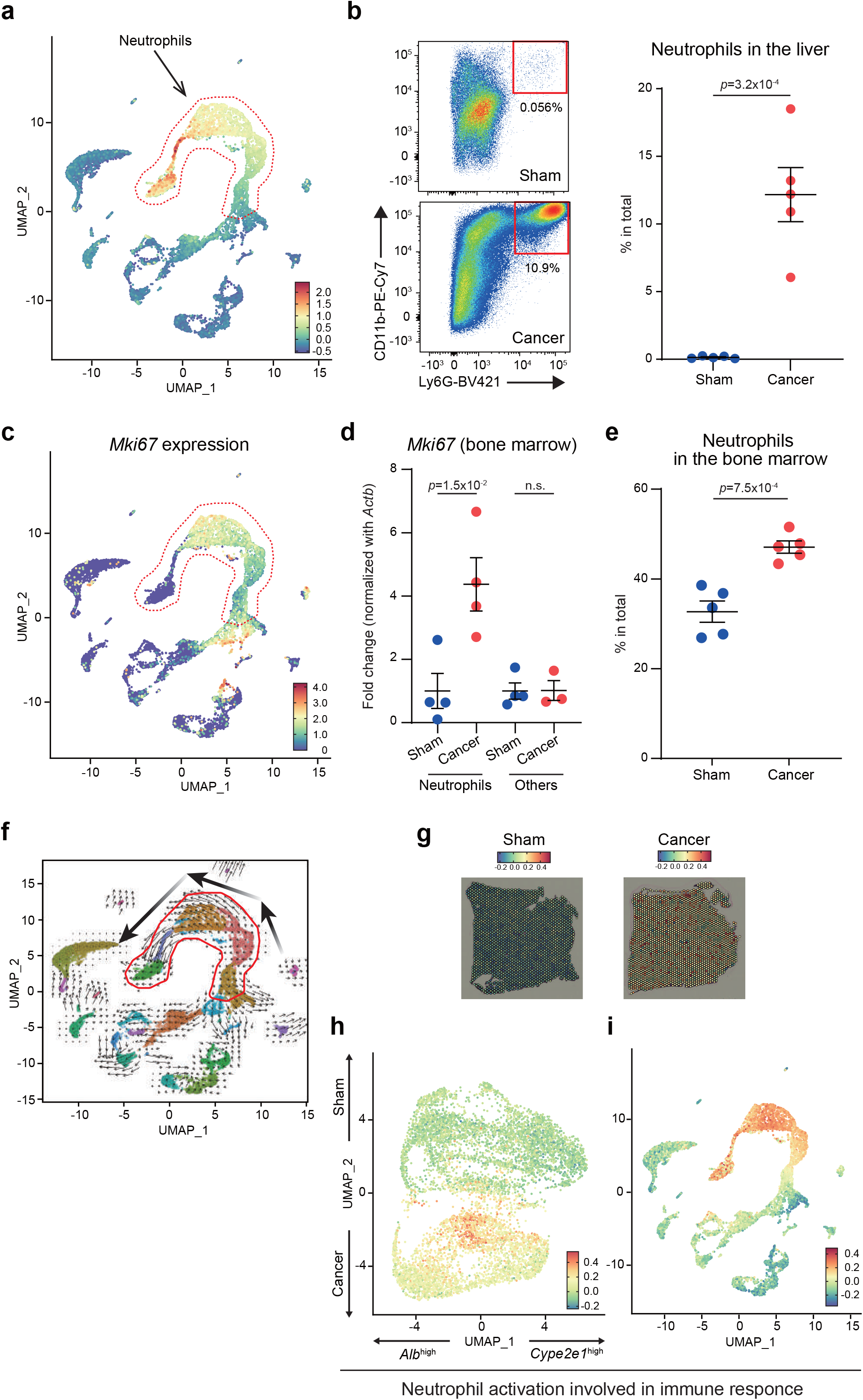
Zonated immune cell activation in the livers of cancer-bearing mice. **(a)** UMAP plot of the scRNA-seq data showing the module score of neutrophil marker genes. **(b)** Flow cytometric analysis of Ly6G^+^CD11b^+^ neutrophils in the livers of sham and 4T1-bearing mice. Representative plots are shown in the left. Data are represented as the mean ± SEM. The *p* value is shown (unpaired two-tailed Student’s *t*-test). *n* = 5. **(c)** UMAP plot of the scRNA-seq data showing the expression of *Mki67.* **(d)** Expression of *Mki67* in the bone marrow of sham and 4T1-bearing mice. Neutrophils and non-neutrophils are MACS-sorted and analyzed by qPCR. Data are represented as the mean ± SEM. The *p* value is shown (unpaired two-tailed Student’s *t*-test). *n* = 4. **(e)** Flow cytometric analysis of Ly6G^+^CD11b^+^ neutrophils in the bone marrow of sham and 4T1-bearing mice. Data are represented as the mean ± SEM. The *p* value is shown (unpaired two-tailed Student’s *t*-test). *n* = 5. **(f)** UMAP plot of the scRNA-seq data showing the predicted dynamics of cells using RNA Velocity. The three big arrows indicate the dynamics of immature neutrophils toward mature neutrophils. **(g-i)** The module scores of genes associated with GO term neutrophil mediated immunity in the Visium slices **(g)** and in a UMAP plot of the same data **(h)**. **(i)** Module scores in the scRNA-seq data, showing high scores predominantly in the neutrophil populations.

### Altered hepatocyte gene expression in zonated manners

As shown in Fig. 2d, there was a general downregulation of genes related to metabolic processes in hepatocytes. This trend was validated by bulk RNA-seq datasets and our previous report^8^, suggesting that breast cancer transplantation repressed metabolism in the liver (Fig. S5). As an example, *Cyp2a22* was down-regulated in the whole liver. Intriguingly, among the repressed processes, we observed a mixture of processes (and genes) that followed zonation patterns, and some that did not. For example, the zonation pattern of genes involved in “xenobiotic catabolic processes” appeared almost completely lost in cancer-bearing samples (Fig. 2d). In contrast, genes involved in “aspartate family amino acid metabolic processes” were generally downregulated in the cancer-bearing samples, yet continued to show a clear zonation pattern (higher expression in the *Alb*^high^ zone; Fig. 2d).

We tried to further support these observations by visualization of Visium spots and UMAP plots. As exemplified by the xenobiotic metabolism pathway, the zonated pattern of this pathway seemed strongly disrupted by 4T1 breast cancers (Fig. 3a). This was evident in the corresponding UMAP plot, where the biased localization of this pathway towards *Cyp2e1^hgil^* spots was no longer detectable (Fig. 3b). Importantly, scRNA-seq datasets validated that cells enriched for this biological pathway were indeed hepatocytes (Fig. 3c). This was a typical pattern of pathways showing “loss of zonation” accompanied by general down-regulation of gene expression. On the other hand, despite being down-regulated, zonation remained relatively intact for “aspartate family amino acid metabolic processes” in *Alb^high^* zones (Fig. 3d-f). We observed a similar trend in other pathways such as the “triglyceride catabolic process” (Fig. 2d and Fig. S6a-c). These results indicated that the effects of breast cancers on liver zonation differ depending on biological pathways.

We also noted that the acute phase response was massively induced in hepatocytes in somewhat zonated manners (Fig. 4a). *Serum amyloid alpha (Saa)* 1-2 were representative of this observation (Fig. S6d, e). These genes were expressed at a very low level in the sham livers but strongly induced by cancer transplantation. Interestingly, this induction occurred in a zonated manner: acute phase response genes were abundantly expressed in *Alb*^high^ zones (Fig. 4b-c), and was driven predominantly by hepatocytes (Fig. 4d). This result together with the repressed metabolism in *Alb*^high^ hepatocytes implicate hepatocytes in switching their focus from metabolism to acute phase response in the presence of breast cancers. These results indicated that breast cancers affected hepatocyte gene expression in zonated manners, which is possibly related to the disrupted liver metabolism observed in the bulk metabolome data we previously reported^4,8^.

### Zonated immune cell activation in the liver

The heatmap shown in Fig. 4a suggested that breast cancers induced immune cell activation and infiltration in the liver. Indeed, the most striking difference between the sham and the cancer-bearing scRNA-seq samples was the appearance of a spectrum of neutrophils in the cancer-bearing liver data (Fig. 5a and Fig. S7). We validated the increase in neutrophils in the liver using flow cytometry (Fig. 5b). Interestingly, these neutrophils were positive for *Mki67,* a marker for cell proliferation (Fig. 5c). This implied that neutrophil proliferation might be enhanced in the bone marrow. To verify this possibility, we used MAgnetic Cell Sorting (MACS) to separate neutrophils and other cell types in bone marrow, and quantified the expression of *Mki67* in both populations by qPCR. We found that *Mki67* was already high in bone marrow neutrophils, supporting our hypothesis (Fig. 5d). Moreover, the neutrophil proportion was elevated in the bone marrow (Fig. 5e). RNA velocity analysis using velocyto.R suggested the existence of a flow between several of the neutrophils clusters (Fig. 5f), in particular from clusters that are exclusively observed in the cancer-bearing samples towards the cluster observed almost exclusively in the sham samples^27^. Interestingly, this flow was correlated with the expression of various neutrophil marker genes such as *Cxcr2* (Fig. S8): *Cxcr2* was strongly expressed in the sham subcluster of neutrophils (i.e., normal mature neutrophils). Other genes such as *Mpo* were strongly expressed in the cancer-specific clusters that were enriched for *Mki67*. These results led to the hypothesis that neutrophils immigrating into the liver of cancer-bearing mice are predominantly immature, differentiating neutrophils, whereas neutrophils in the livers of sham mice are mature neutrophils.

Our data suggested a biased (i.e., non-uniform) localization of these neutrophils in the livers of 4T1 cancer-bearing mice. Although the zonation pattern was hard to spot by eye in the Visium slices, the pattern was clear in the UMAP plot, showing especially high signals in the *Alb*^inter^ zone of the cancer-bearing liver (Fig. 5g-i). This pattern was also seen in other immune-related pathways as shown in Fig. 4a. These results suggested an influx of neutrophils into the livers of cancer-bearing animals, preferentially to regions of the liver that are not high in *Alb* expression, but rather between the portal and central veins. Indeed, several marker genes of neutrophils had particularly high expression levels at this subset of spots (Fig. S9). These data were in line with the heatmap showing the activation of the immune system in zonated manners (Fig. 4a; see for example “neutrophil activation involved in immune response”), suggesting that neutrophil distribution in the livers of cancer-bearing mice was not uniform.

Although we initially focused on neutrophils, we also found the activation of other cell types. We found that macrophages were activated in zonated manners, most likely representing activation of liver resident macrophages (i.e., Kupffer cells) (Fig. S10a-c). Macrophage spots were observed from the *Alb^inter^* to *Alb^high^* zones, a pattern distinguishable from neutrophils (Fig. 5h). The increase of basophils was also notable in scRNA-seq and flow cytometry (Fig. 2f and Fig. S10d, e). Basophils were not present in the sham controls in the scRNA-seq datasets (Fig. S7). Using flow cytometry, we detected an increase in basophils in the liver whereas they were reduced in the bone marrow. These results suggested the enhanced influx of basophils from the bone marrow to the liver (Fig. S10d, e). Unlike neutrophils, it appeared that basophils exhibited neither *in situ* differentiation nor zonated localization. Given their roles in expressing various unique cytokines, these innate immune cells might contribute to the altered spatial transcriptomics landscape of the livers in various distinct manners.

Together, our results suggest that the induced gene expression in cancer-bearing mice could be attributed to at least two different processes; the influx of immune cells (especially neutrophils) and the response of liver-resident cells (especially hepatocytes) to distal cancer and/or the influx of immune cells.

## Discussion

### Disrupted liver zonation in cancer-bearing mice

Although it is known that solid cancers remotely disrupt liver homeostasis, it has been unclear whether this disruption accompanies abnormalities in liver zonation. In the current study, we addressed this question by combining spatial transcriptomics and single-cell transcriptomics, finding that a murine breast cancer model disorganizes liver transcriptome in zonated manners.

Our analyses integrating spatial transcriptomics and single-cell transcriptomics allowed us to categorize breast cancer-induced gene expression changes in the bulk livers according to liver zonation. We found that some gene expression changes were coincident with loss of zonation, as exemplified by the xenobiotic catabolic process in the *Cyp2a1^high^* zone (Fig. 3a-c). On the other hand, zonation of the aspartate family amino acid metabolism and the triglyceride catabolic process was relatively intact in the *Alb^high^* zone (Fig. 3d-f and Fig. S6a-c). In addition, acute phase protein genes *Saa1* and *Saa2* showed an intriguing pattern of activation: their activation was not uniform within the liver. Rather, *Saa1* and *Saa2* were induced in the *Alb^high^* zone (i.e., gain of zonation), suggesting a zonation-distinct gene expression mechanism underlying acute phase response (Fig. 4 and Fig. S6d, e). These results indicate that the effects of breast cancers on liver zonation were distinct according to biological pathways, revealing previously unrecognized complexity of cancer-induced gene expression changes in the liver. It remains unclear how these pathway-specific changes in zonation occurred. Further detailed investigation addressing this issue is critical to deepening our knowledge of how solid cancers and other diseases disrupt liver homeostasis.

The increased influx of immune cells to various peripheral organs is a hallmark of host pathophysiology in cancers^1,10,28^. This is also the case for the liver, and many studies pointed out the massive infiltration and activation of particularly innate immune cells in the cancer-bearing condition^2–5,8,28^. Our data detected various zonated patterns of immune cell activation in the liver (Fig. 5 and Fig. S9–10). We found that potentially immature neutrophils infiltrated the liver in the *Alb^inter^* zone. It seems that they were gradually changing their transcriptome status within the liver, possibly implicating differentiation *in situ* (Fig. 5f). We also captured the zonated activation of liver-resident macrophages without such *in situ* transcriptomic changes (Fig. S10a-c). Basophils followed neither the neutrophil pattern nor the macrophage pattern (Fig. S10d, e). It appeared that basophils migrated from the bone marrow to the liver, forming a single cluster. Collectively, our analyses revealed intriguing, distinct patterns of influx and activation of various innate immune cells to the liver, which we assume are important to understand the basis of the altered liver zonation and disrupted liver homeostasis in the cancer-bearing condition.

How these unique patterns are generated and how such altered immune cell dynamics affect other cell types especially hepatocytes are important questions to be answered.

Our data revealed other zonated patterns of gene expression in the livers of cancer-bearing mice whose significance is relatively unclear to us at present (Fig. S11). The transcytosis pathway was much more active in *Alb*^high^ zones rather than in *Cyp2e1*^high^ zones (Fig. 4a and Fig. S11a-c). 4T1 cancer transplantation elevated the expression of genes involved in this pathway, potentially enhancing transcytosis in endothelial cells nearby portal veins. In an interesting contrast, the transepithelial transport appeared zonated towards *Cyp2e1*^high^ zones and activated in the presence of 4T1 breast cancers (Fig. 4a and Fig. S11d-f). A previous study reported immune cell zonation orchestrated by liver sinusoidal endothelial cells^23^. In this report, the authors suggested that liver sinusoidal endothelial cells sense the microbiome, accordingly modulating immune cell localization. The complex interaction among different cell types and stimuli might underlie the altered zonation of immune cells and endothelial cells in our experimental settings. Addressing these zonation patterns in detail is also an important next step.

### Disrupted liver zonation in various disease conditions

Our study is currently limited to a murine breast cancer model, but a disease-dependent disruption in liver zonation could be general^15^. For example, pathogenesis such as non-alcoholic fatty liver disease (NAFLD), non-alcoholic steatohepatitis (NASH), and liver cirrhosis develops in a somewhat zonated manner. In detail, these develop with steatosis and inflammation in the pericentral region^29,30^. In an intriguing contrast, various liver injuries begin periportally^31–33^. These indicate that each liver pathology has a distinct pattern of disrupted liver zonation, which possibly underlies the nature of such pathology. Obtaining spatial transcriptomics datasets from various disease conditions will be useful to advance our understanding of various liver diseases in light of alterations in zonated gene expression.

Given the nature of cancer transplantation models, disease progression in our experimental settings could be considered extreme. Compared to other pathologies described above, which initiate locally in the specific region of the liver, the altered zonation in our model was consistent throughout the spatial transcriptome sections. These results led to an interpretation that the effects of cancer transplantation on liver zonation were strong, reflecting the terminal state of diseases. We imagine that disruption in zonated gene expression might begin in more region-specific manners in the livers of e.g., metastatic cancer patients. The lack of human data is a limitation in our study, and future investigation of liver zonation in human cancer patients is critical to address the hypothesis we described. Yet, we believe that the current datasets from a murine breast cancer model will be a basis for gaining insights into the terminal state of liver abnormalities in the cancer-bearing condition.

In summary, we demonstrate uniquely altered patterns of zonated gene expression in the liver. Our study highlights the strengths of the combination of spatial transcriptomics and single-cell transcriptomics to understand cancer’s adverse effects on the liver and to reveal complex interactions among various cell types in disease conditions.

## Methods

### Mice

All animal experiment protocols were approved by the Animal Care and Use committee of Kyoto University. Wild-type female mice (8-week-old) purchased from Japan SLC Inc. (Hamamatsu, Japan) were housed in a 12-hour light/dark paradigm with food ((CE-2, CLEA Japan, Inc., Tokyo) and water available *ad libitum.* No blinding was done in the experiments described in this study. WT female mice were purchased from Japan SLC Inc. (Hamamatsu, Japan).

### Cell line

4T1 mouse breast cancer cell line^8^ was cultured in RPMI1640 (nacalai tesque, Kyoto, Japan) supplemented with 10% fetal bovine serum (nacalai tesque) and 1% penicillin/streptomycin (nacalai tesque) in a 5% CO2 tissue culture incubator at 37°C.

### Spatial transcriptomics

Cancer transplantation was performed as described previously^8^. 2.5×10^6^ 4T1 cells were inoculated subcutaneously into the right flank of 8-10-week-old BALB/c females and sacrificed 14 days after transplantation and livers were harvested. The obtained specimens were embedded in OCT Compound (Sakura Finetek Japan, Tokyo, Japan) and frozen samples embedded in OCT compound were sectioned at a thickness of 10 μm (Leica CM3050S). We utilized the 10x Genomics Visium Spatial Gene Expression Solution for spatial transcriptomics. Libraries for Visium were prepared according to the Visium Spatial Gene Expression User Guide (https://www.10xgenomics.com/). Mouse livers were mounted for 4 stages (two sham samples and two 4T1 cancer-bearing samples) with one stage per capture area on a 10x Visium gene expression slide containing four capture areas with 5000 barcoded RNA spots printed per capture area. Spatially barcoded cDNA libraries were built using the 10x Genomics Visium Spatial Gene Expression 3’Library Construction V1 Kit. The optimal permeabilization time for 10 μm thick liver sections was determined to be 6 minutes using 10x Genomics Visium Tissue Optimization Kit. H&E staining of the liver tissue sections was imaged using a light microscope (Keyence BZ-X800, Keyence Ltd. HQ & Laboratories, Osaka, Japan), and the images were stitched and processed using Keyence BZ-X800 Analyzer software.

The library was sequenced using the NovaSeq 6000 system (Illumina) according to the manufacturer’s instructions. Sequencing was carried out using a 28/90bp paired-end configuration, which is sufficient to align confidentially to the transcriptome. Raw FASTQ files and histology images aligned to the mouse reference genome provided by 10x Genomics (mm10 Reference-2020-A) using the Space Ranger 1.2.1 pipeline to derive a feature spot-barcode expression matrix (10x Genomics).

### Spatial transcriptomics data analysis

#### Data processing

Spatial transcriptome data, including UMI counts and spot coordinates, were analyzed using the R Seurat package (version 4.0.0)^34^ The four images and their data were processed into Seurat objects using the Load10X_Spatial function and merged into a single Seurat object. This resulted in a dataset of 7,758 spots and 32,285 genes. Data were processed using a standard Seurat workflow (SCTransform function). We conducted batch effect reduction between the four slices using the SelectIntegrationFeatures, PrepSCTIntegration, FindIntegrationAnchors, and IntegrateData functions, followed by dimensionality reduction using PCA and UMAP (based on 30 principal components).

#### Prediction of spatially variable genes

We predicted spatially variable genes (SVGs) using singleCellHaystack (version 0.3.4)^26^. The haystack_highD function detects genes whose expression is distributed in a non-random pattern inside the input space (= here the 2D spatial coordinates of the spots). For each of the 4 slices, the inputs to the haystack_highD function were 1) the spatial coordinates of the spots and 2) the detection data of each gene at each spot. Genes were defined as “detected” if their signal in a spot was higher than the median signal in the sample. Additional input parameters to haystack_highD were grid.points = 500, and scale = FALSE. To avoid biases by damaged cells, spots with a high proportion of mitochondrial UMIs (top 10%) were excluded from this analysis.

#### Analysis of zonation of genes sets and their module scores

Using the R package msigdbr (version 7.5.1) we defined sets of genes associated with Gene Ontology (GO) biological process functional annotations. After filtering out GO terms associated with < 20 or > 250 genes present in our data, we retained 2,898 GO terms and their associated genes. For each of these sets, we calculated module scores using Seurat’s AddModuleScore function in both the spots of the spatial transcriptomics data as well as in the individual cells of our scRNA-seq data. These module scores reflect a weighted average expression level of the gene set.

To characterize zonation in the patterns of activity of gene sets, we divided the Visium spots of each image into 3 sets of equal size: those with the highest *Alb* expression (*Alb*^high^), those with the highest *Cyp2e1* expression (*Cyp2e1*^high^), and the remaining one-third (with intermediate *Alb* and *Cyp2e1* expression. Next, we compared the module scores of each GO term in the *Alb*^high^ zone versus the *Cyp2e1*^high^ zone using the Wilcoxon Rank Sum test (R function wilcox.test). Differences in the mean module scores of the *Alb*^high^ zone versus the *Cyp2e1*^high^ zone (X-axis) and the p-value of the Wilcoxon Rank Sum test (log_10_; Y-axis) were visualized using a volcano plot (Fig. 1b).

To characterize the zonation patterns at a higher resolution, we first defined *Alb*^high^ spots as the top 10% of spots with the highest *Alb* expression in each image. In the Visium platform, spots are organized in a hexagonal pattern. Each spot has 6 closest neighbors (at a distance of 100 μm center-to-center), and 6 secondary neighbors (at a distance of about 173 μm center-to-center). For each image, we defined *Cyp2e1*^high^ spots and their closest and secondary neighbors in the same way. Next, for each GO term, we calculated the mean of the module scores in the *Alb*^high^ and *Cyp2e1*^high^ spots and their neighbors (Table S3). The 200 GO terms with the largest ranges of values were picked up for visualization using heatmaps (Fig. 2d and 4a).

### Single-cell RNA-seq

Cancer transplantation was performed as described in the section “spatial transcriptomics.” The obtained livers were minced and dissociated in RPMI1640 containing DNase I (100 μg/ml: Roche, Switzerland) and collagenase IV (1 mg/ml: Worthington Biochemical Corporation, NJ, USA). Red blood cells (RBCs) were lysed with RBC lysis buffer (BD Bioscience, NJ, USA). The obtained cells were stained with anti-CD45 antibody (Clone: 30-F11 (BVD421)), anti-CD19 antibody (Clone: 1D3), and anti-TCRβ antibody (Clone: H57-597) for 30 minutes at 4°C. Then we sorted and mixed 20,000 CD45^-^ cells that include hepatocytes, 6,000 CD45^+^CD19^+^TCRβ^-^ cells (i.e., B cells), 6,000 CD45^+^CD19^-^TCRβ^+^ cells (i.e., T cells), and 20,000 CD45^+^CD19^-^TCRβ^-^ cells (e.g., neutrophils) using FACS Aria (BD Bioscience, NJ, USA). The obtained mixture of cells was subjected to single-cell transcriptomics.

The single-cell RNA-seq library was constructed by using the Chromium Controller and Chromium Single Cell 3’Reagent Kits v3.1 (10x Genomics) following the standard manufacturer’s protocols. Cells sorted from fresh live mouse liver cells were washed with PBS, then pipetted through a 40-μm filter to remove cell doublets and contamination. Cell viability was confirmed by trypan blue staining. The collected single-cell suspension was immediately loaded onto the 10x Chromium controller to recover 10000 cells, followed by library construction. The library was sequenced using the NovaSeq 6000 system (Illumina). Sequencing was carried out using a 28/90bp paired-end configuration. After sequencing analysis, the obtained fastq files were mapped to the reference genome provided by 10x Genomics (refdata-cellranger-mm10-3.0.0) and quantified read count using the Cell Ranger ver 3.1.0 count pipeline (10x Genomics) with the default parameters.

### Single-cell RNA-seq data analysis

The UMI counts for the four samples were analyzed using the R Seurat package (version 4.0.0)^34^. Initial exploratory analysis revealed that one of the sham samples contained a large population of cells with a relatively high fraction of mitochondrial reads, which formed a separate cluster of cells after dimensionality reduction. To avoid biases caused by these cells, we applied a more stringent filter for this sample, removing cells with more than 5% mitochondrial UMIs. For the remaining 3 samples, we removed cells with more than 20% mitochondrial UMIs. In addition, for all samples, we removed cells with fewer than 200 detected genes. This resulted in a dataset with 11,085 cells (sham 1: 364; sham 2: 2,660; cancer-bearing 1: 4,769; cancer-bearing 2: 3,302) and 19,355 genes.

Data were processed and subjected to dimensionality reduction using a standard Seurat workflow, including normalized, selection of 2000 variable features, scaling, principal component analysis (PCA), and UMAP using the first 20 PCs. The clustering of cells was done using Seurat’s FindNeighbors function (using 20 PCs) and FindClusters function (using resolution 0.5), resulting in 23 clusters.

We assigned a cell type to each cluster after inspection of the expression patterns of a selection of known marker genes (Table S3).

RNA Velocity analysis was performed using velocyto.py (version 0.17.17), and the R packages velocyto.R (version 0.6), and SeuratWrappers (version 0.3.0) (https://github.com/satijalab/seurat-wrappers)^27^. First, we ran velocyto.py on each of the 4 scRNA-seq samples using option run10x, the mm10 gene annotation file, and a repeat annotation file (option -m) obtained from the UCSC Genome Browser^35^. Resulting.loom files were processed and visualized using functions ReadVelocity, RunVelocity, and show.velocity.on.embedding.cor.

### Bulk RNA-seq data analysis

We used DESeq2 (version 1.28.1) to normalize the read count per gene for all 8 samples (4 sham and 4 cancer-bearing)^8^. DEGs between sham and cancer-bearing samples were predicted using default parameters.

### Flow cytometry

The livers were harvested from sham and 4T1 cancer-bearing animals 14 days after transplantation. Red blood cells (RBCs) were lysed with RBC lysis buffer (BD Bioscience). The obtained cells were then stained with the following antibodies (BioLegend, CA, USA) for 30 minutes at 4°C: anti-CD45 antibody (Clone: 30-F11), anti-CD19 antibody (Clone: 6D5), anti-TCRβ antibody (Clone: H57-597), anti-CD11b antibody (Clone: M1/70), anti-Ly6G antibody (Clone: 1A8), anti-FCεR1a antibody (Clone: MAR-1), anti-CD117 antibody (Clone: 2B8), anti-CD123 (Clone: 5B11), anti-CD49b antibody (Clone: DX5), anti-CD3 antibody (Clone: 17A2), anti-CD11c antibody (Clone: N418), anti-TER-119 antibody or anti-CD34 antibody (Clone: HM34). Non-viable cells were stained with Propidium Iodide Solution (PI) and gated out. Data were acquired using FACS Aria (BD Bioscience) and analyzed using FlowJo software (v10.8.1, BD Biosciences).

### MACS

The mouse hind limb femur and tibia from both legs of sham and 4T1-bearing mice were collected in RPMI1640 media (nacalai tesque) in a 60 mm dish. The femur and tibia were transferred to a new dish with RPMI1640 medium after associated muscles and connective tissues were removed. The femur and tibia were cut and perfused using a syringe needle. The suspension containing immune cells was transferred to a clean tube. Neutrophils and the other cell types were isolated from the suspension using the MACS Neutrophil isolation kit (Miltenyi Biotec, 130-097-658) according to the manufacturer’s instructions. The obtained cells were then subjected to RNA isolation, cDNA synthesis, and qPCR analysis.

### qPCR

The collected murine livers were crushed in liquid nitrogen and homogenized with Trizol reagent (Thermo Fisher Scientific) as described previously^8^. Total RNA was extracted from the homogenized supernatant using RNeasy Mini Kit (Qiagen, Venlo, Netherlands) according to the manufacturer’s instruction and reverse-transcribed with the aid of Transcriptor First Strand cDNA synthesis kit (Roche, Switzerland). qPCR experiments were performed using the StepOnePlus qPCR system (Applied Biosystems, CA, USA) and SYBR Green Master Mix (Roche, Basal, Switzerland). *Actb* was used as an internal control. The primers used in these experiments are:

*Mki67* (Forward: AAAAGTAAGTGTGCCTGCCC,

Reverse: GGAAAGTACGGAGCCTGTAT),

*Actb* (Forward: CGGTTCCGATGCCCTGAGGCTCTT,

Reverse: CGTCACACTTCATGATGGAATTGA)

### Data availability

The Visium and scRNA-seq datasets in this study are available in DNA Databank of Japan (DDBJ) under the accession numbers of DRA014802 and DRA009332, respectively. The codes used in this study will be provided upon request.

## Acknowledgement

We thank Dr. Diego Diez (Osaka University) for helpful discussions and advice. This work was supported by JSPS KAKENHI (16H06279, 18K15409, 18H04810, 20H03451, 20H04842, and 22H04925; S.K, and 20K06609; A.V.), JST FOREST (20351876; S.K), JST Moonshot (JPMJMS2011-61; S.K), Caravel, Co., Ltd (S.K), Ono Medical Research Foundation (S.K), Takeda Science Foundation, The Uehara memorial foundation (S.K), Chubei Ito foundation (S.K) and Japan Foundation for Applied Enzymology (S.K).

We thank Yuri Kawaoka for drawing the cartoons in Fig.2e.

## Competing interests

The authors declare no competing interests exist in this study.

## Author contribution

A.V. performed and supervised data analyses, constructed figures, and wrote the manuscript. R.M, and R.K performed in vivo experiments, analyzed data, and constructed figures. M.O performed sequencing data acquisition, analyzed data, and made a substantial contribution in the discussion regarding the relevance of our work to human cancers. K.M, Y.K, and H.K supported in obtaining Visium slices and flow cytometry analyses. A.S supported initial data analyses. M.S. and Y.T. made substantial contribution to the conception of this work. Y.S. supervised Visium and scRNA-seq experiments and data analysis. S.K. conceived and supervised this study, managed the collaboration in this study, analyzed data, and wrote the manuscript.

---

**Supplementary Figure S1:**
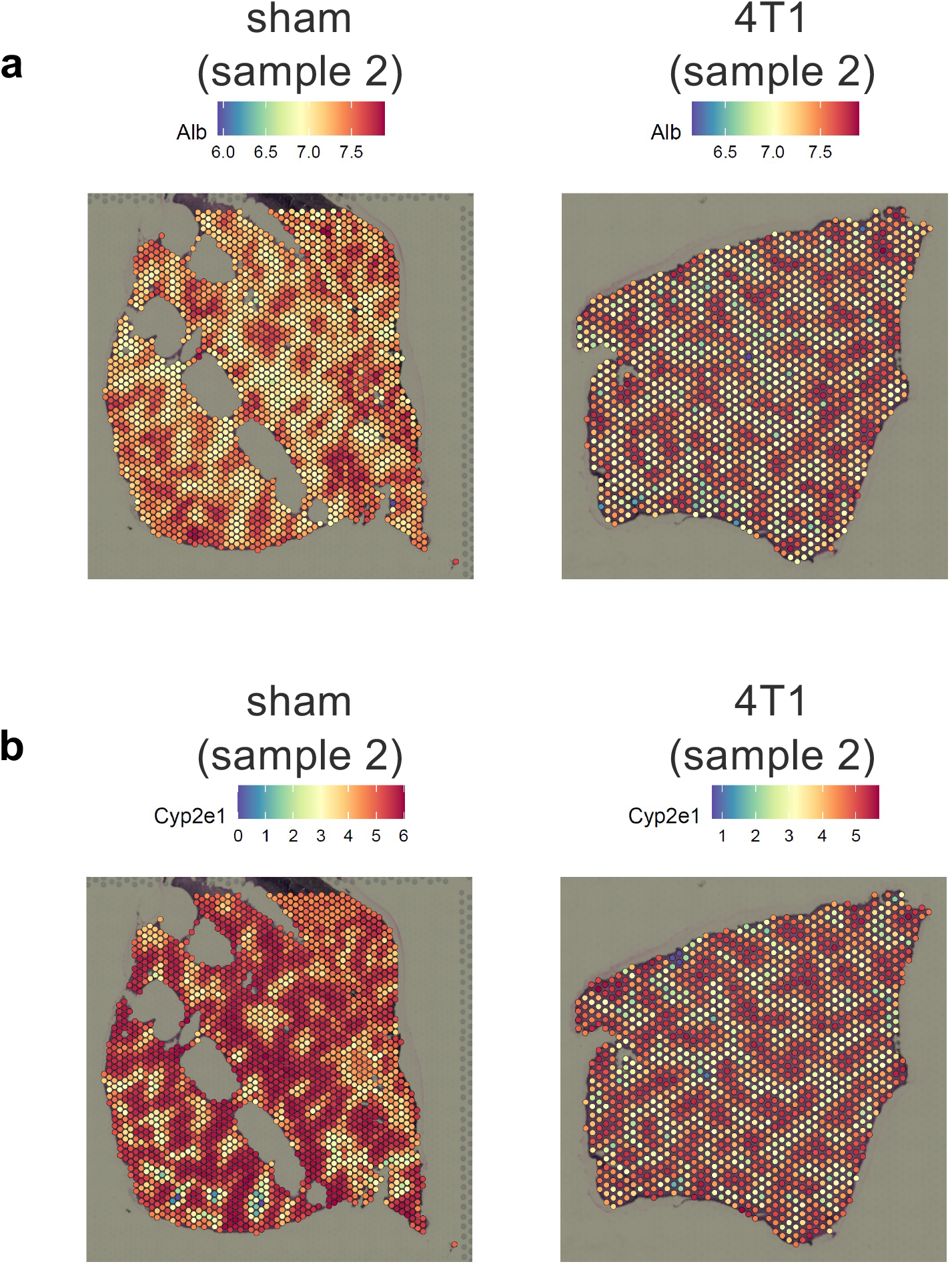
Expression of *Alb* and *Cyp2e1* in three Visium liver samples. Expression of *Alb* **(a)** and *Cyp2e1* **(b)** is shown in the second sham sample (left), and the second 4T1 cancer-bearing sample (right). The corresponding figures for the first sham sample are shown in main **Fig. 1a** and for the first cancer-bearing sample in **Fig. 2a**.

**Supplementary Figure S2:**
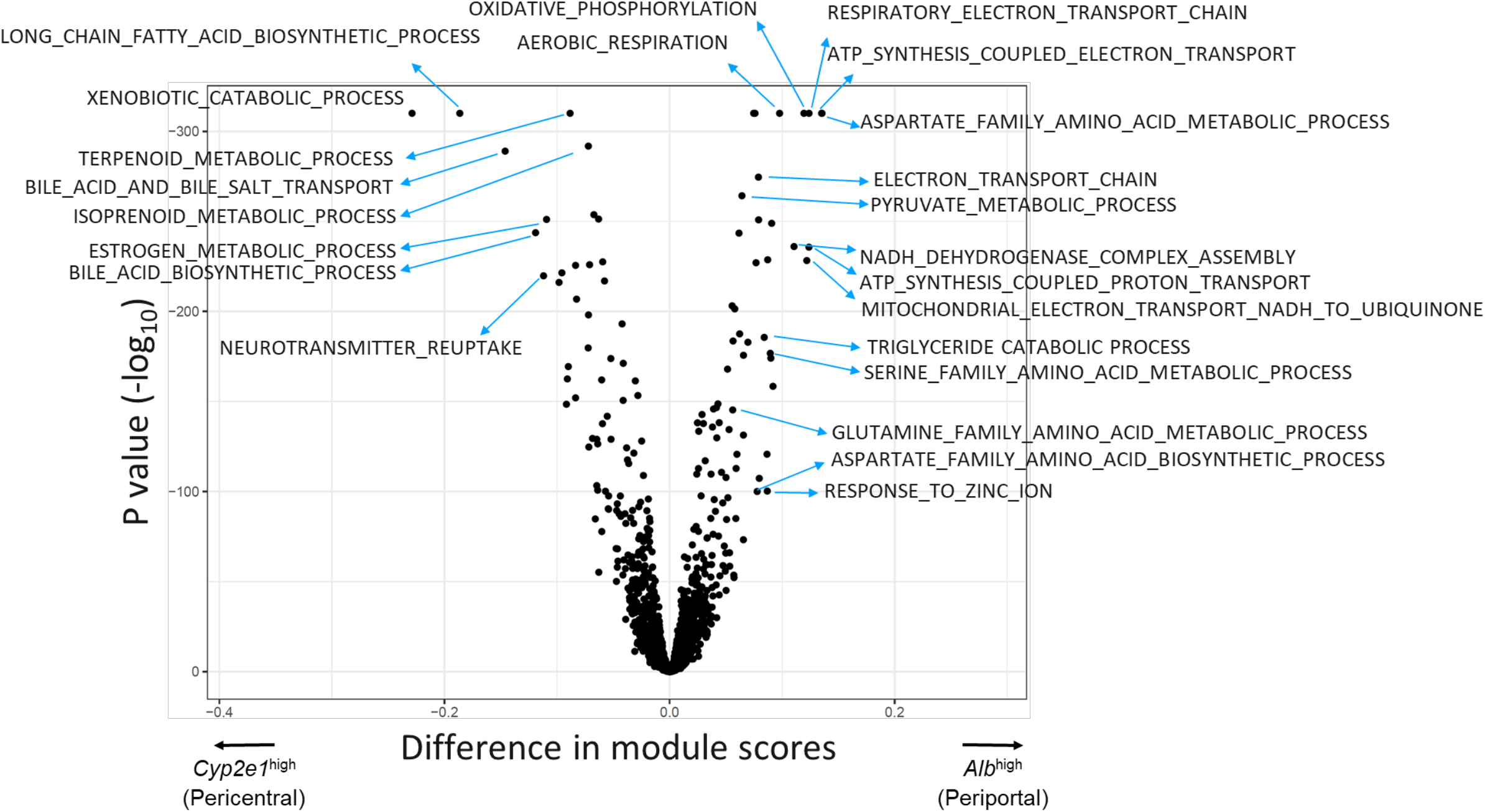
Volcano plot of the module scores of 2,898 GO terms. This figure supports Fig. 2b in the main manuscript. The X-axis shows the difference in module scores between *Alb^high^* and *Cyp2e1^high^* zones. The Y-axis shows the *p* values (−log_10_) of a Wilcoxon Rank Sum test.

**Supplementary Figure S3:**
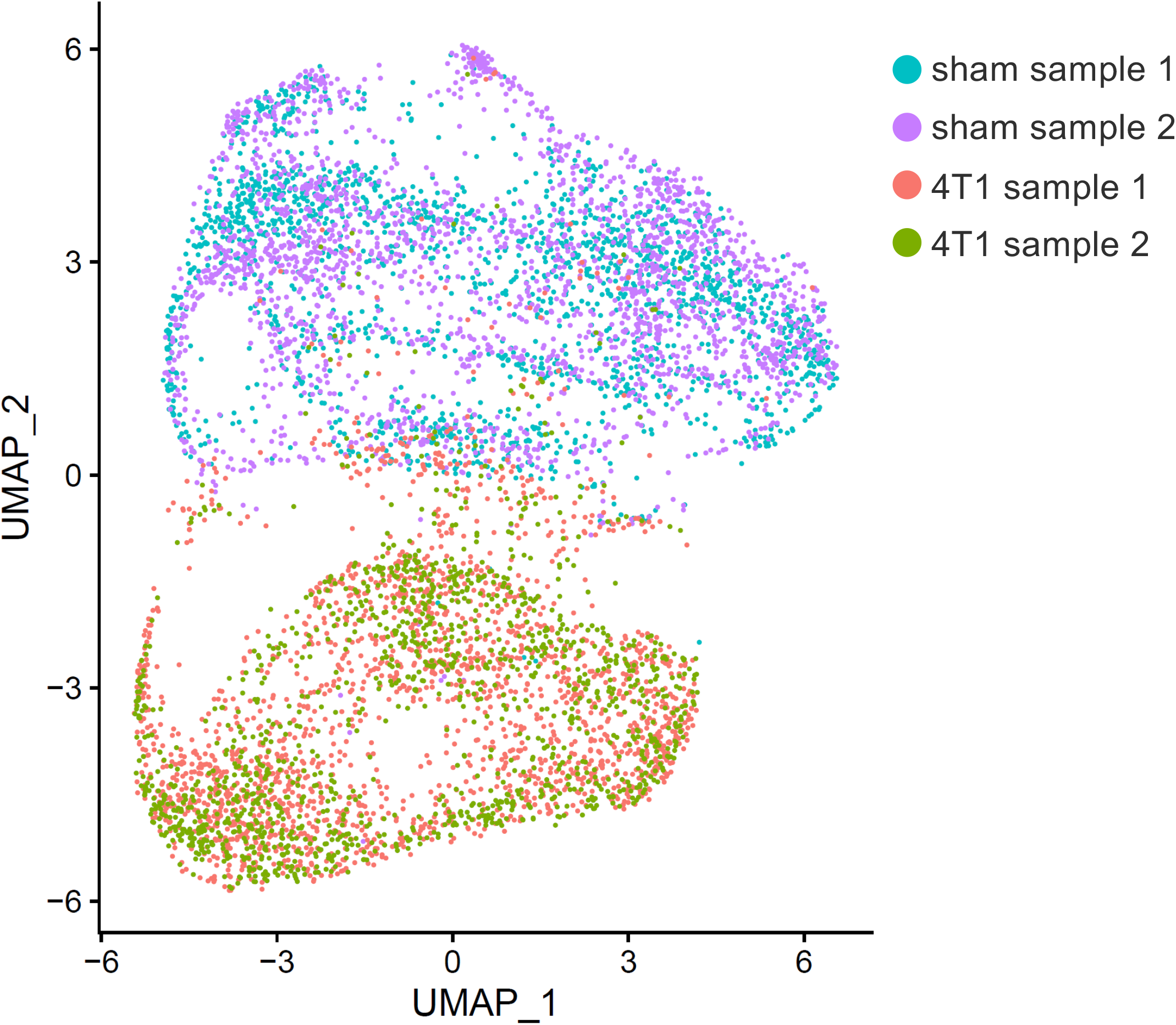
UMAP plot of the spots of the four Visium samples. Colors represent the samples (two sham and two 4T1 cancer-bearing samples). Even though spots of the two sham samples and the two cancer-bearing samples are well-mixed, there is a clear separation between the sham and cancer-bearing samples, suggesting a clear difference in gene expression.

**Supplementary Figure S4:**
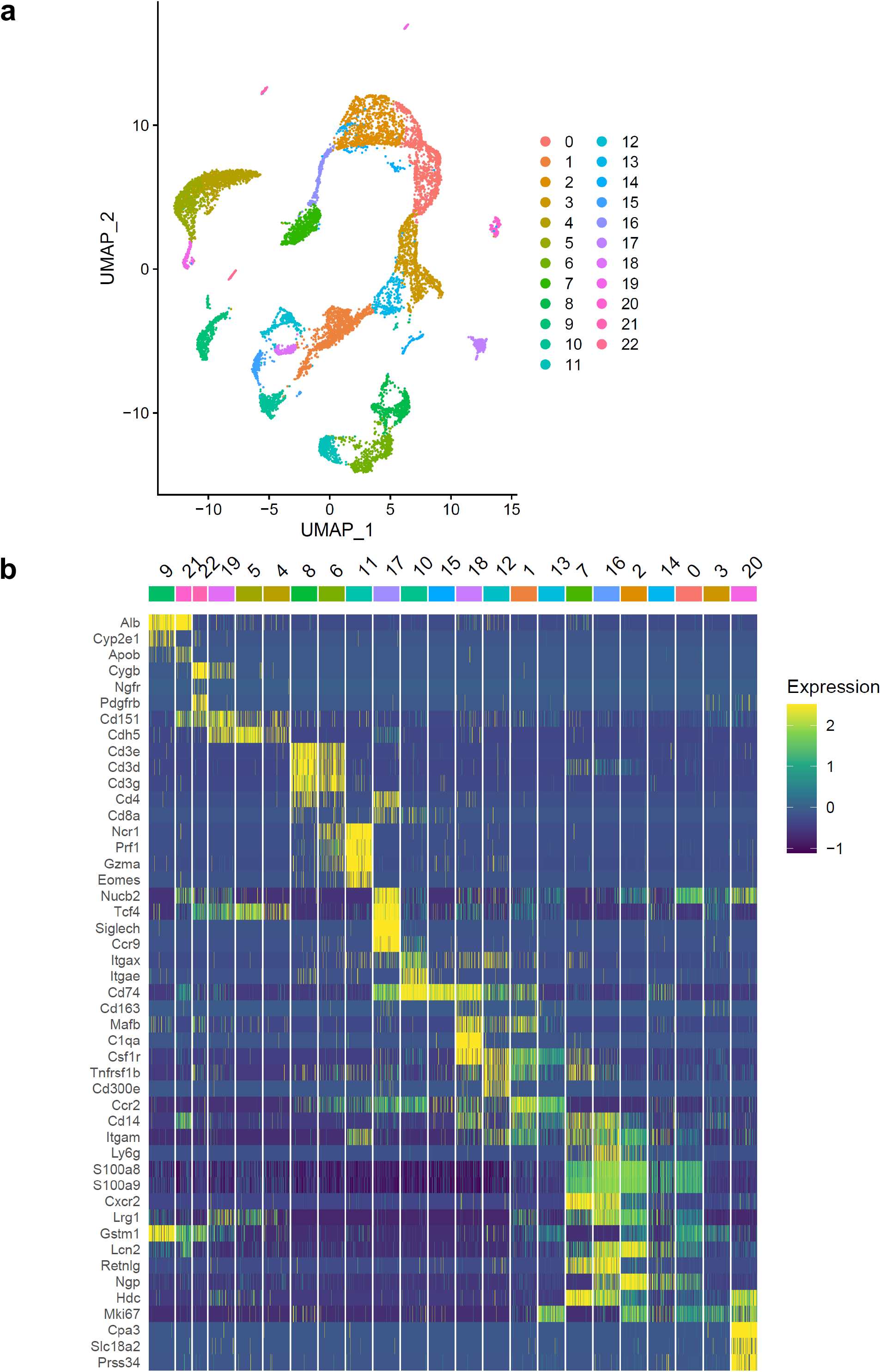
Clusters and cell type annotations of the scRNA-seq data. **(a)** UMAP plot of the scRNA-seq data showing the 23 detected clusters. **(b)** Scaled expression patterns of a selection of cell type marker genes used for making cell type assignments (see **Fig. 2f**).

**Supplementary Figure S5:**
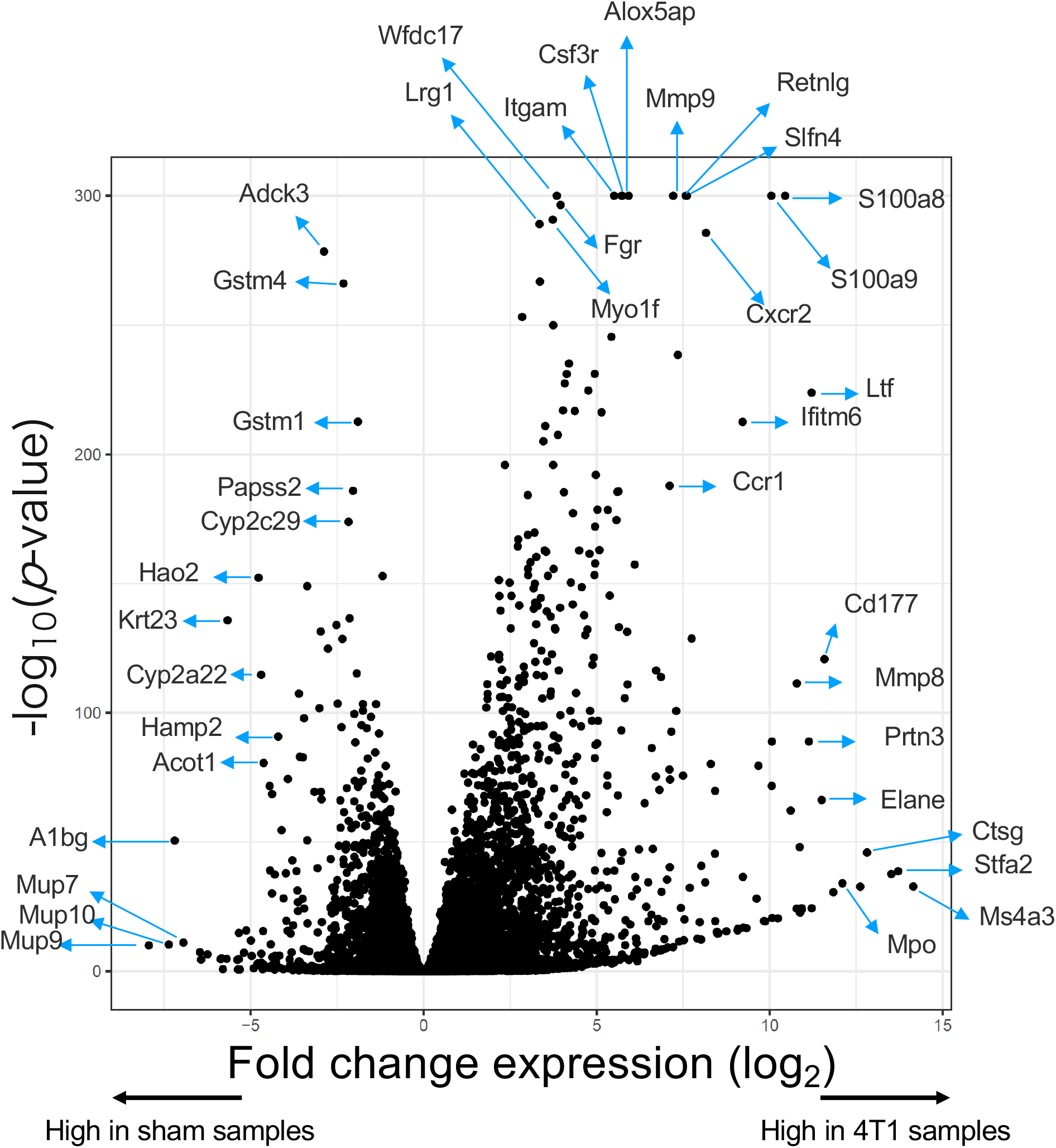
Volcano plot of bulk RNA-seq data showing changes in gene expression in livers of sham and 4T1 cancer bearing mice. The X-axis represents fold changes (log_2_ values) and the Y-axis represents *p* values (-log_10_ values) based on a comparison using DESeq2 between four sham and four cancer-bearing samples. Genes of interest are indicated.

**Supplementary Figure S6:**
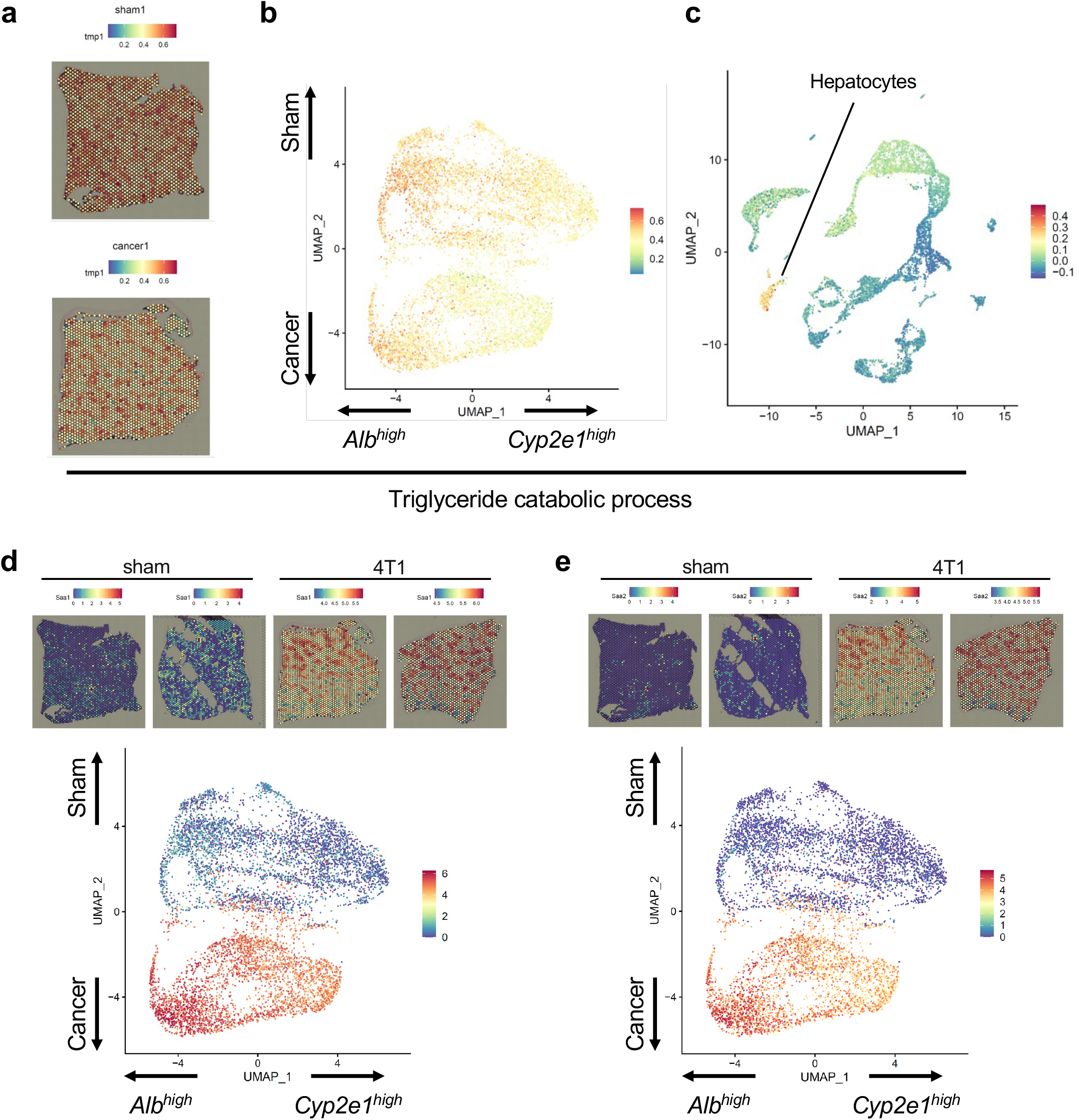
Additional examples of the effects of 4T1 breast cancers on liver zonation. **(a-c)** Genes associated with triglyceride catabolic process are relatively unaffected by 4T1 breast cancers. **(a)** module scores of genes associated with triglyceride catabolic processes in one of the sham and cancer Visium samples. **(b)** The same module scores in a UMAP representation of the Visium data. **(c)** Module scores in the scRNA-seq data, showing high scores predominantly in the hepatocyte cluster. **(d-e)** Gene expression patterns of *Saa1* **(d)** and *Saa2* **(e)** in the Visium data. Both genes are strongly induced in the livers of caner-bearing mice, especially in the periportal (i.e.*Alb*^high^) part of the liver.

**Supplementary Figure S7:**
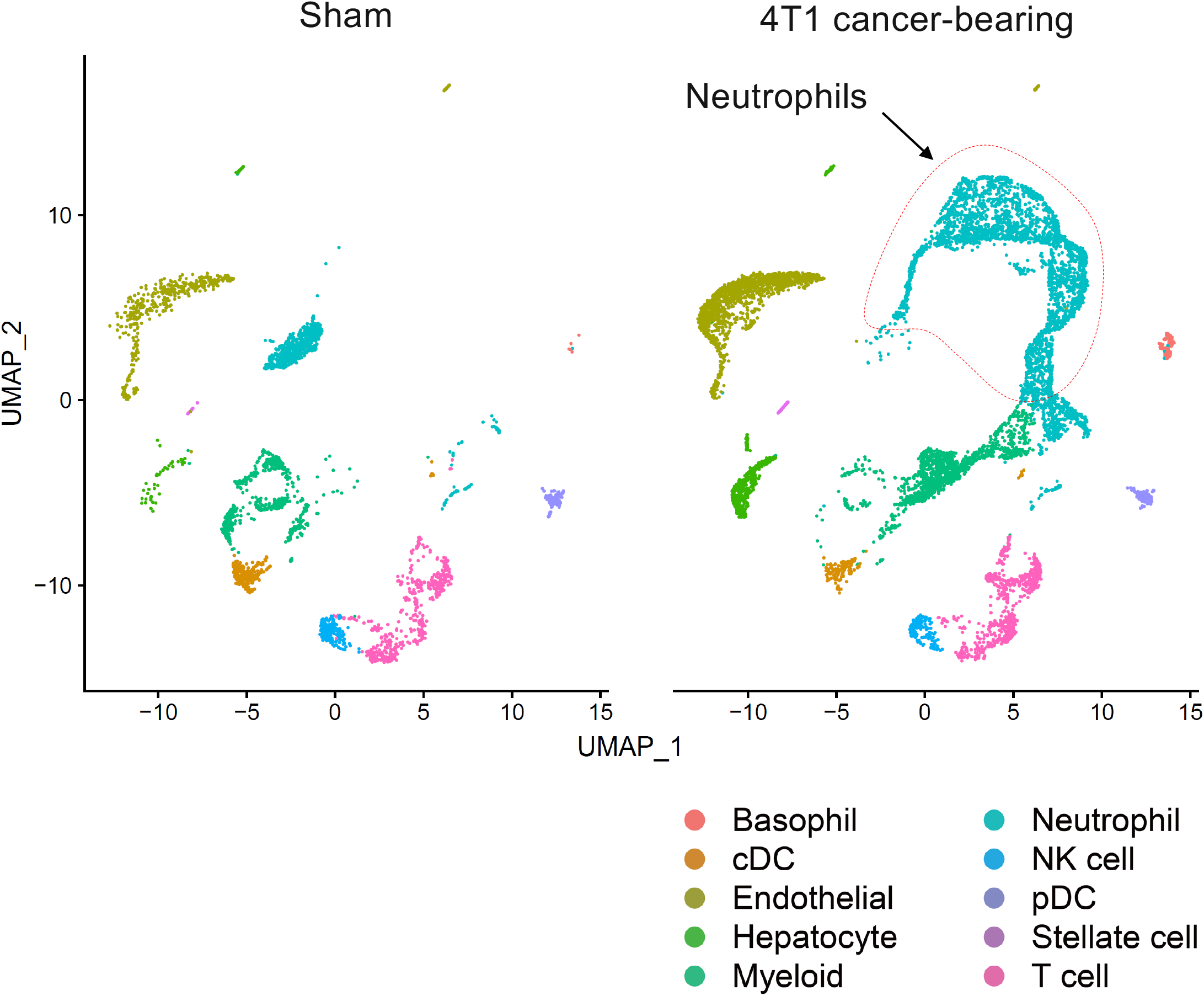
UMAP plot of the scRNA-seq dataset separated by condition. Colors represent cell type annotations. The subpopulation of neutrophils infiltrating into the liver of 4T1 cancer-bearing mice is indicated.

**Supplementary Figure S8:**
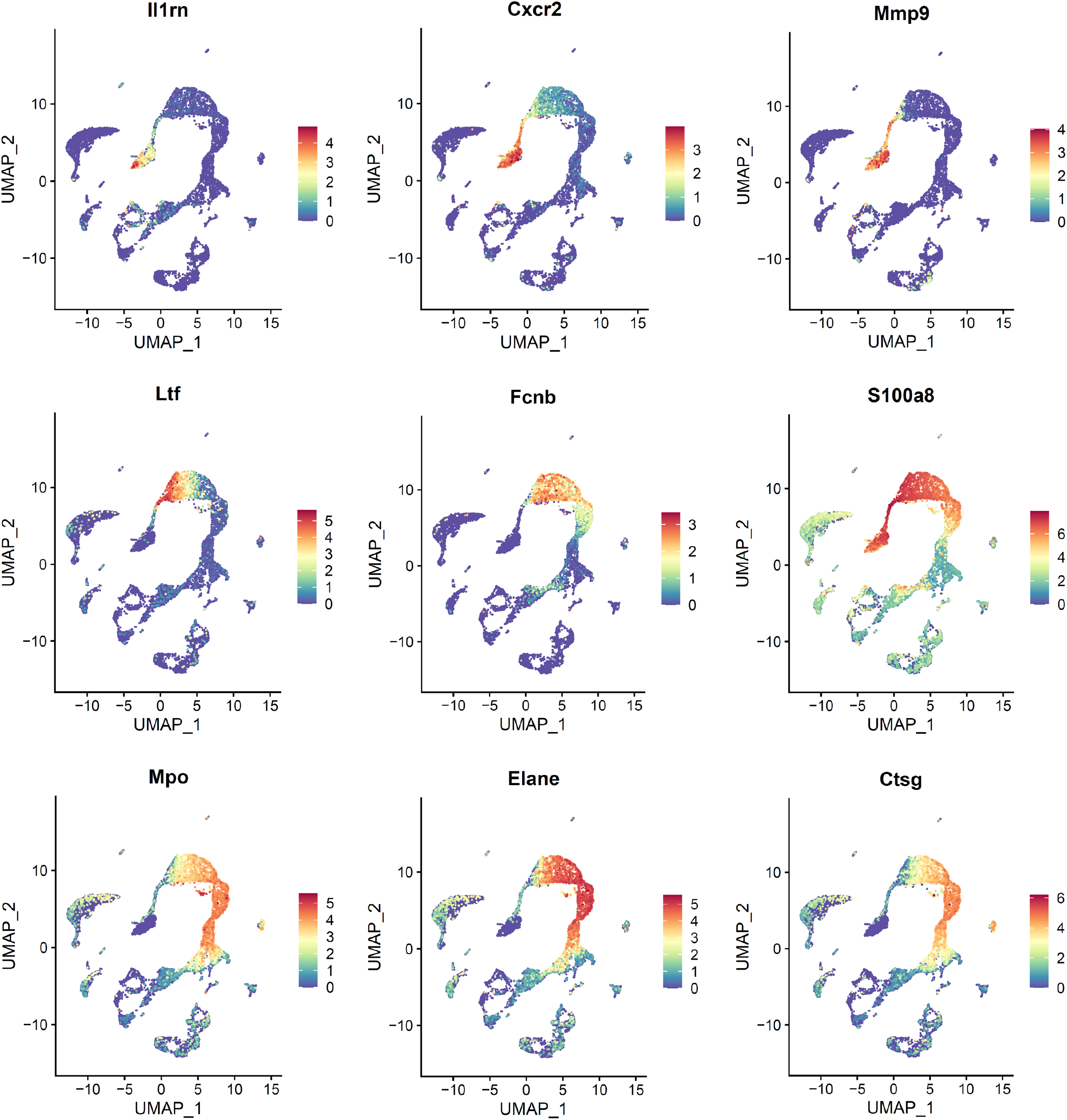
UMAP plots of the scRNA-seq data showing the variety of expression patterns between several neutrophil marker genes.

**Supplementary Figure S9:**
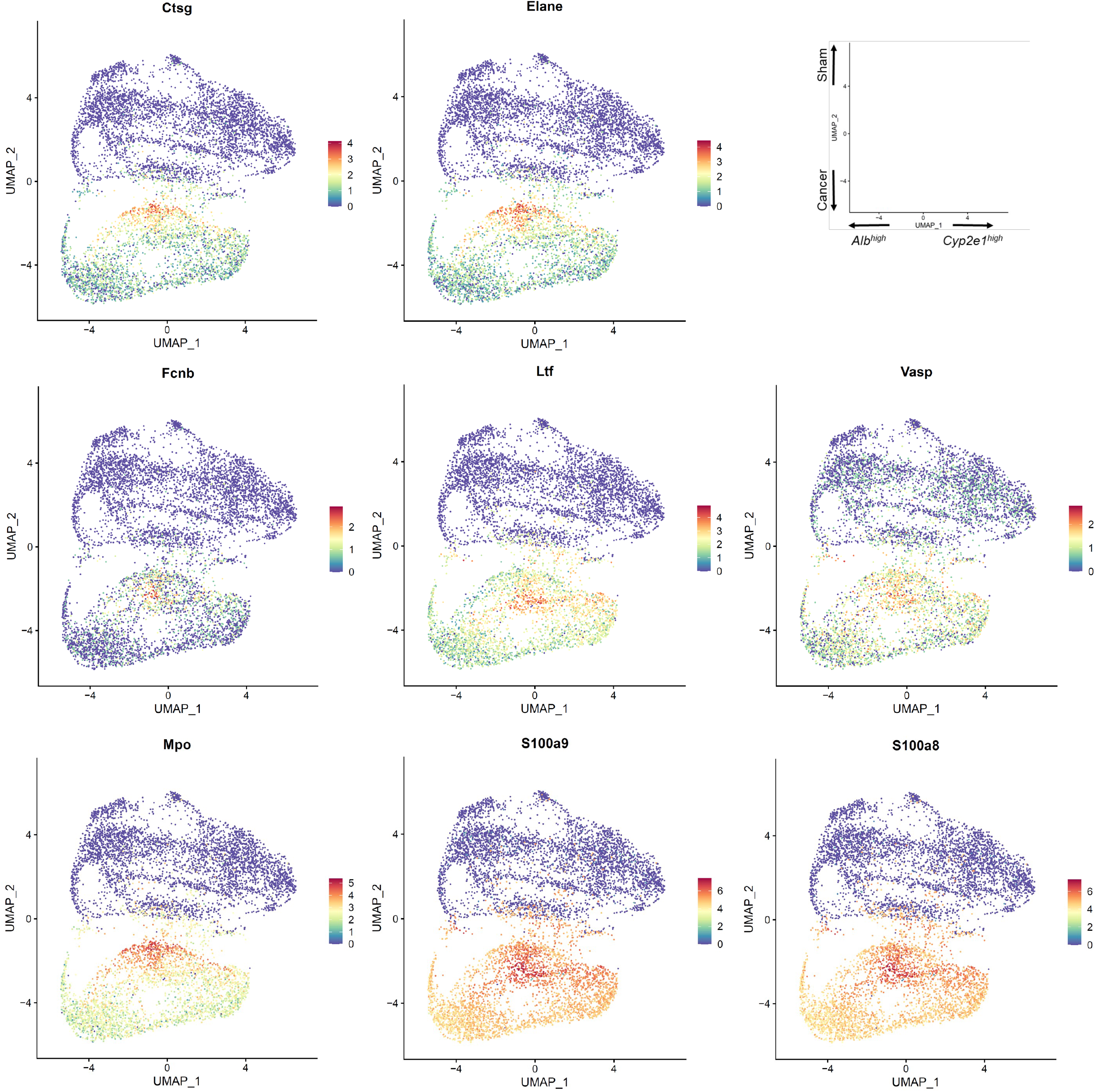
UMAP plots of the Visium datasets showing the expression patterns of several neutrophil marker genes.

**Supplementary Figure S10:**
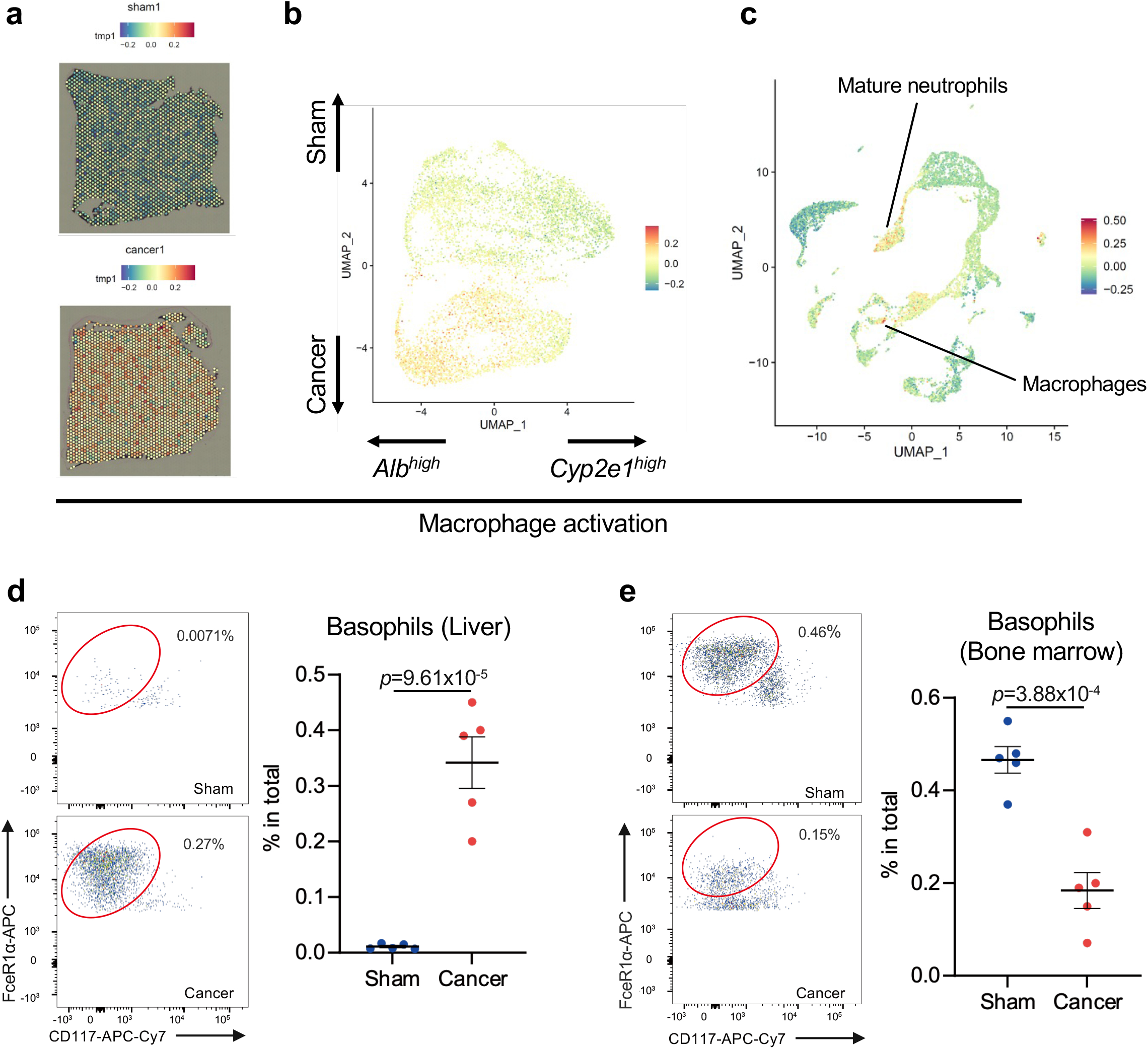
Immune cell activation in the livers of cancer-bearing mice. **(a-c)** Genes associated with macrophage activation are induced by 4T1 breast cancers. **(a)** module scores of genes associated with macrophage activation in one of the sham and cancer Visium samples. **(b)** The same module scores in a UMAP representation of the Visium data. **(c)** Module scores in the scRNA-seq data, showing high scores predominantly in a subset of neutrophils and macrophage cells. **(d-e)** Flow cytometric analysis of FceR1α^+^CD117^-^basophils in **(d)** the livers and **(e)** bone marrows of sham and 4T1-bearing mice. Representative plots are shown in the left. Data are represented as the mean ± SEM. The *p* value is shown (unpaired two-tailed Student’s *t*-test). *n* =5.

**Supplementary Figure S11:**
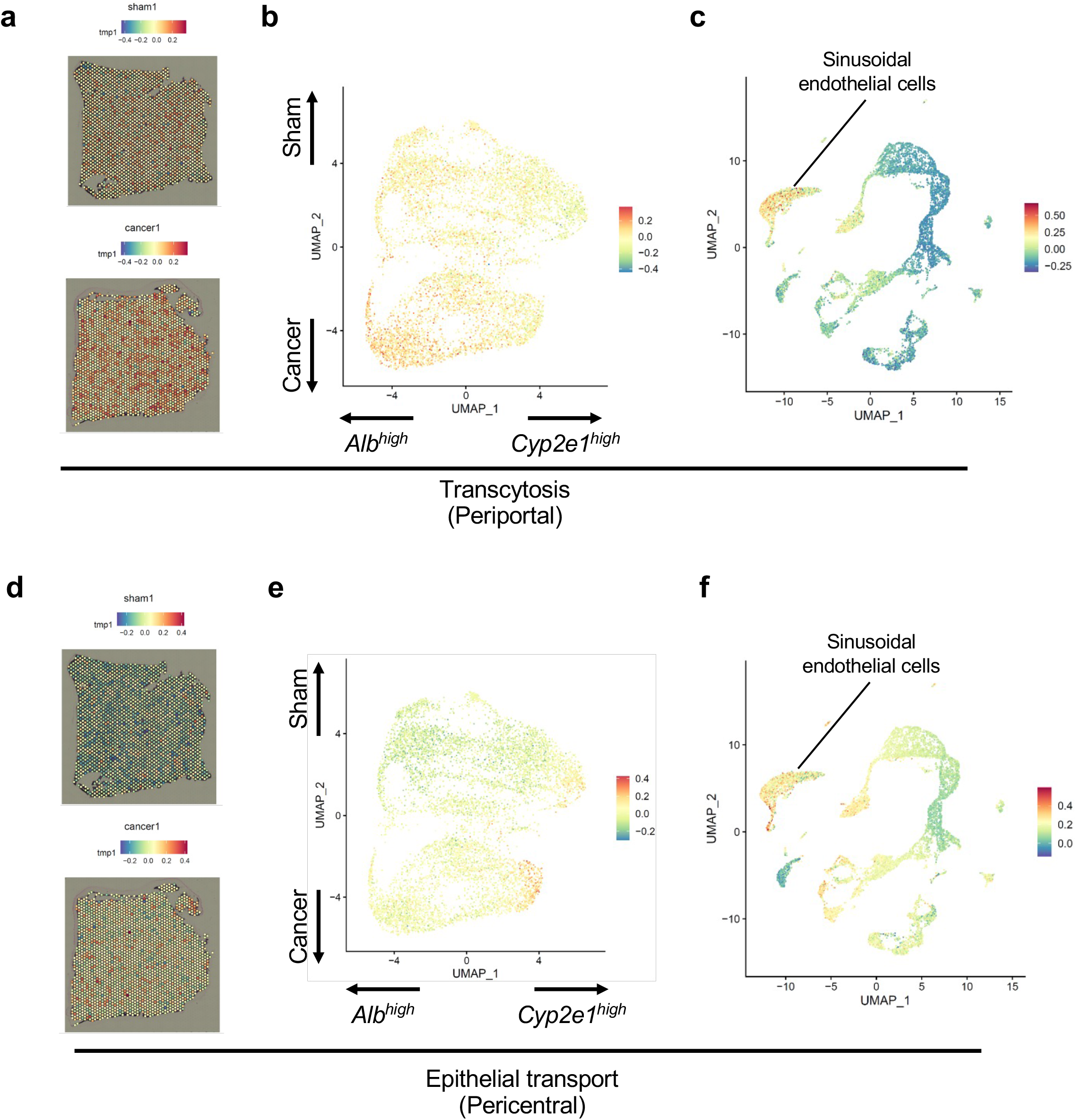
Zonated expression patterns of genes involved in transcytosis and epithelial transport. **(a-c)** Zonated expression patterns of genes associated with transcytosis. **(a)** module scores of genes associated transcytosis in one of the sham and cancer Visium samples. **(b)** The same module scores in a UMAP representation of the Visium data. **(c)** Module scores in the scRNA-seq data, showing high scores predominantly in the cluster of endothelial cells. **(d-f)** Similar plots showing the zonated expression patterns of genes involved in epithelial transport.

**Supplementary Figure S12:**
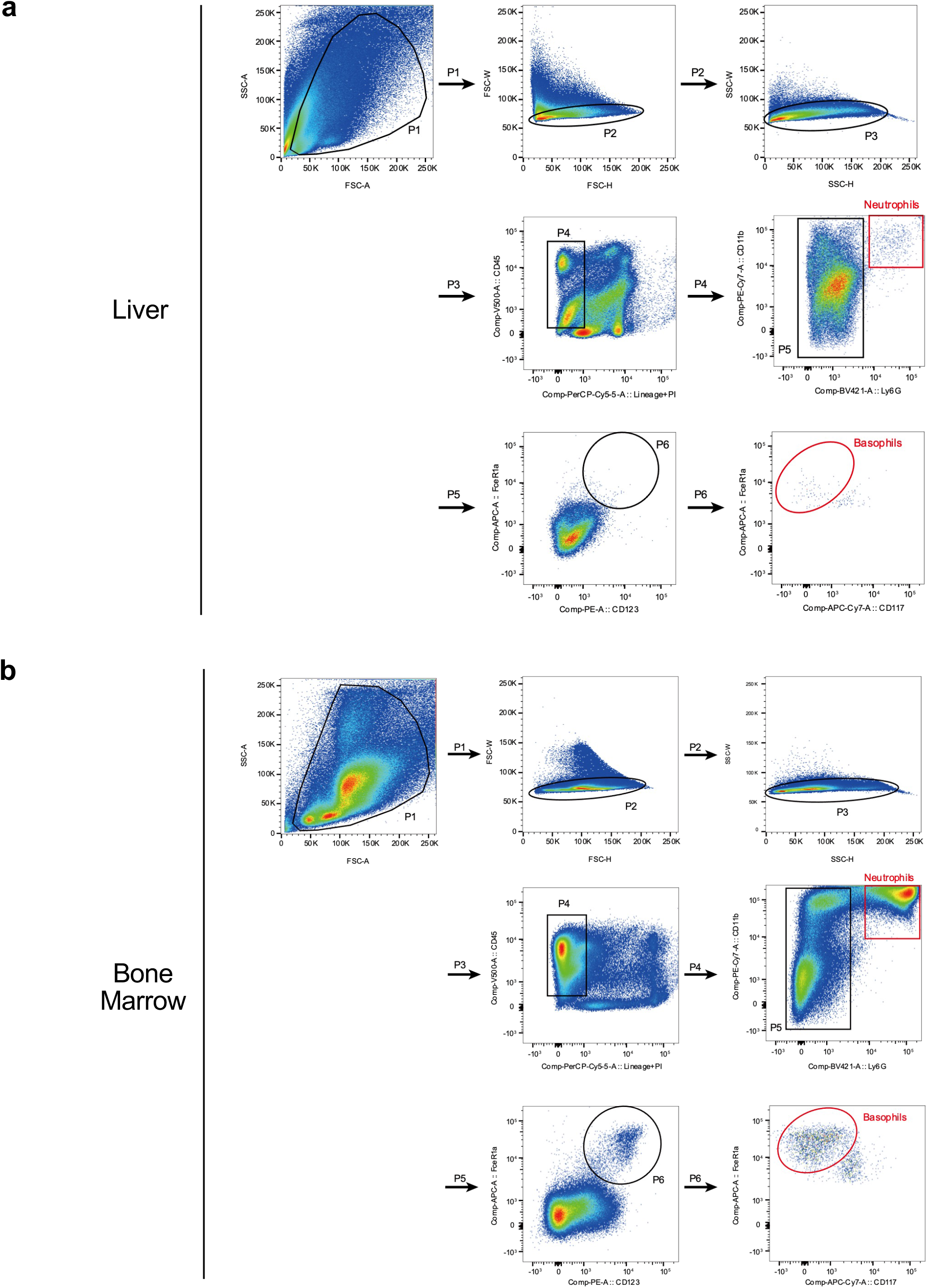
Gating strategies for neutrophils and basophils. (a) Liver. (b) Bone marrow.

## Notes

### Competing Interest Statement

The authors have declared no competing interest.

